# Identifying Emerging Phenomenon in Plant Long Temporal Phenotyping Experiments

**DOI:** 10.1101/454686

**Authors:** Jiajie Peng, Junya Lu, Donghee Hoh, Ayesha S Dina, Xuequn Shang, David M Kramer, Jin Chen

## Abstract

**The rapid improvement of phenotyping capability, accuracy, and throughput have greatly increased the volume and diversity of phenomics data. A remaining challenge is an efficient way to identify phenotypic patterns to improve our understanding of the quantitative variation of complex phenotypes, and to attribute gene functions. To address this challenge, we developed a new algorithm to identify emerging phenomena from large-scale temporal plant phenotyping experiments. An emerging phenomenon is defined as a group of genotypes who exhibit a coherent phenotype pattern during a relatively short time. Emerging phenomena are highly transient and diverse, and are dependent in complex ways on both environmental conditions and development. Identifying emerging phenomena may help biologists to examine potential relationships among phenotypes and genotypes in a genetically diverse population and to associate such relationships with the change of environments or development. We present an emerging phenomenon identification tool called Temporal Emerging Phenomenon Finder (TEP-Finder). Using large-scale longitudinal phenomics data as input, TEP-Finder first encodes the complicated phenotypic patterns into a dynamic phenotype network. Then, emerging phenomena in different temporal scales are identified from dynamic phenotype network using a maximal clique based approach. Meanwhile, a directed acyclic network of emerging phenomena is composed to model the relationships among the emerging phenomena. The experiment that compares TEP-Finder with two state-of-art algorithms shows that the emerging phenomena identified by TEP-Finder are more functionally specific, robust, and biologically significant. The source code, manual, and sample data of TEP-Finder are all available at: http://phenomics.uky.edu/TEP-Finder/.**

## I. Introduction

Biomedical studies have been ushered into a new era by the rapid development of large-scale genotyping and phenotyping technologies [2], [8], [11], [13], [17]. Recent studies demonstrate that by integrating both phenomics and genomics, we can better understand organism behaviors and identify new genes that govern phenotypes and response to the varying environments [7], [9], [29]. More specifically, by analyzing large-scale plant photosynthetic phenotype data, researchers can identify complex aggregate phenotypic traits, and explore the processes or genetic components that control a trait and the essential conditions under which the trait emerge [15], [31].

The main computational challenge in omics data analysis arises from its unsupervised nature. It is generally believed that the **emerging phenomena** among multiple phenotypes measured across several genotypes (e.g., gene knockouts) reveals, to a great extent, the common regulatory roles of the knocked out genes in the biological system. An emerging phenomenon refers to a phenotypic pattern that multiple genotypes have correlated phenotype values during a serial of continuous time points [8]. A sample emerging phenomenon in plant photosynthesis phenotype data is shown in Figure 1. The experiment was done under the fluctuating light conditions (between 0 and 1000*μmolm*^−2^*s*^−1^). Five selected genotypes (*P*_1_…*P*_5_) were measured using three photosynthetic phenotypes, namely photosynthetic system II activity (Φ_*II*_), photoprotection (*q*_*ESV*_), and photoinhibition (*q*_*I*_). The relative phenotype values were calculated by comparing each genotype with the reference (col-0) using logged fold change. The shadowed area indicates an emerging phenomenon of the five plants between 12:30 and 14:30, during which, all the five genotypes have similar phenotype values.

**Figure 1.**
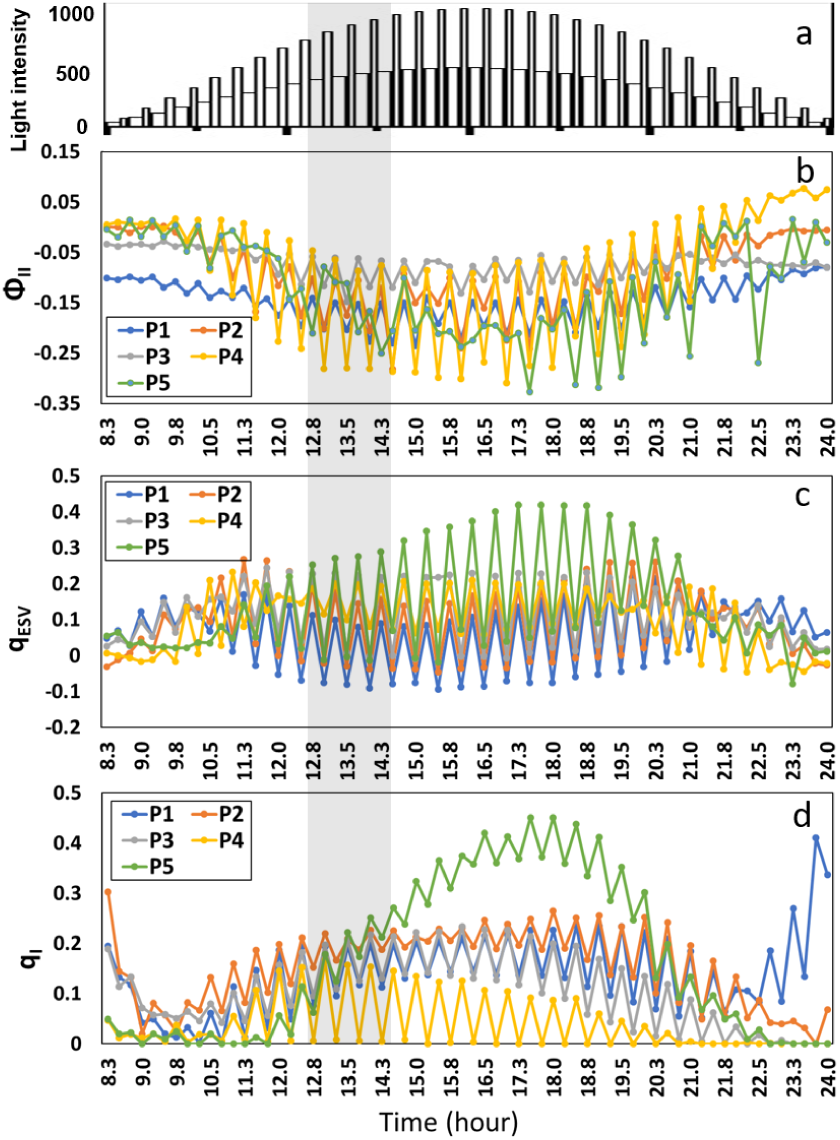
A sample emerging phenomenon (shadowed area) identified in a plant photosynthesis phenotyping experiment under fluctuating light conditions (a). In the experiment, five genotypes (chloroplast-targeted single mutant lines of *Arabidopsis thaliana*) were measured using three photosynthetic phenotypes (Φ_*II*_, *q*_*ESV*_, and *qI*). In the shadowed area (b,c,d), all the genotypes have similar phenotype values.

Emerging phenomena is universal in phenotyping experiments *esp*. under dynamic environmental conditions. They are highly transient and diverse, dependent in complex ways on both environmental conditions and development [8]. Revealing emerging phenomena is vital towards the identification of meaningful differences in biological function among genotypes, which may help biologists to examine potential phenome-genome relationships in a genetically diverse population and to associate such relationships with the change of environments or development. It is, however, *unclear* what specific patterns biomedical researchers should look for given the complexity of the biological system and its responses to environmental perturbations. Besides, the large variance in phenotypes, due to the biodiversity and the variance in environmental distribution, adds more challenges to the already difficult task [8], [11], [33].

To address this challenge, we propose a new tool called **Temporal Emerging Phenomenon Finder** (TEP-Finder) as the first approach to capture emerging phenomena with various temporal scales and arbitrary phenotype variation shapes (see Figure 2). TEP-Finder automatically transforms large-scale phenomics data into emerging phenomenon patterns, thus facilitates the translation of information into knowledge. TEP-Finder has two phases. First, TEP-Finder encodes phenotype-based relationships into a dynamic network using nonparametric clustering and generates seeds. It then identifies all the emerging phenomena in different temporal scales and constructs a directed acyclic network of emerging phenomena. To demonstrate the effectiveness of TEP-Finder, we applied TEP-Finder on a large-scale plant photosynthesis phenotyping experiment, and the results show that TEP-Finder can reliably and accurately identify high quality emerging phenomena from data. Comparing with the existing models, TEP-Finder has the following advantages:

**Figure 2.**
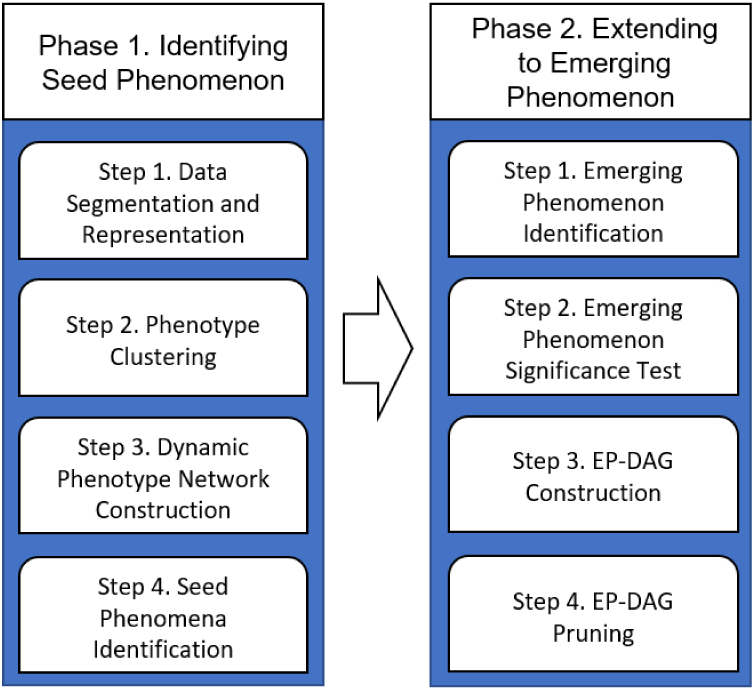
The workflow of TEP-Finder. Given the temporal phenomics data, it identifies significant seed phenomena in every time frame; by expanding each seed to longer time frames, it discovers emerging phenomena that appear and disappear subject to the change of environments or development; the relationships among all the emerging phenomena are modeled by a directed acyclic network called EP-DAG.

- TEP-Finder is the first approach to capture emerging phenomena with diverse scales systematically;
- TEP-Finder constructs a network of emerging phenomena to provides a graph-based representation of the complex hierarchy of emerging phenomena;
- TEP-Finder successfully discovers emerging phenomena in an Arabidopsis photosynthetic phenotyping experimental data with high biological significance.

## II. Background

An emerging phenomenon is defined as a group of genotypes who has a pattern of correlated phenotypes in a serial of continuous time points [8]. In the literature, given a set of predefined patterns, the minimal genotype contributor set can be identified using existing data mining techniques such as association rule mining [16], [32] or subspace trajectory clustering [1], [26]. However, given the unsupervised nature, most emerging phenomena are not pre-definable. To our knowledge, there is no existing algorithm exactly designed for emerging phenomena identification. Tools, such as DHAC and NPM [12], [19], may be slightly modified to achieve the goal. Here we discuss two existing approaches with additional steps adopted for emerging phenomenon discovery.

DHAC models how a network change with time [19]. Assuming that the edges in a network are conditionally independent given group membership, DHAC uses a probabilistic model to translate a hierarchical stochastic block to the dynamic domain, thus clustering a time-evolving network based on the observations at several specific time points. The rationale is that any node in a network cluster at a specific time point should be influenced by clusters at nearby time points. DHAC can be employed to group genotypes by matching clusters across multiple time points with additional steps that transform longitudinal phenomics data into a dynamic network (called DHAC+). However, to facilitate dynamic network clustering, DHAC considers global features on all temporal points rather than local features. Subsequently, the DHAC-based method cannot identify emerging phenomenon at different temporal scales.

NPM is a non-parametric clustering method that can simultaneously cluster subjects with arbitrary cluster shapes [12]. NPM represents the phenotypes of each genotype in a serial of continuous time points as a cloud of points. Each point of the cloud corresponds to a vector in the sequential phenotype measurements taken for the genotype. Two similar shapes of clouds represent that two genotypes have a coherent phenotype pattern in a given time frame. Note that NPM is more advantageous than the Pearson correlation on the identification of a set of genotypes with coherent phenomics data. It is because Pearson correlation requires all the variables to follow a normal distribution, which is not always held for the phenomics data, while NPM does not make any assumption about the underlying data distribution and thus is particularly suitable for phenomics data analysis. NPM can be employed to identify emerging phenomena by applying it repeatedly on every time frame of a longitudinal phenomics dataset (called NPM+). However, it is difficult to pre-define the time scale of emerging phenomena or to identify the relationships between overlapped emerging phenomena. Furthermore, NPM is not a deterministic method so that the results are dependent on the initialization and the selection of anchor points.

The unmet needs to effectively identify high-quality emerging phenomena necessitates the development of tools that can automatically transform large-scale phenomics data into emerging phenomenon patterns, thus facilitate the translation of information into knowledge. To precisely identify emerging phenomena with different temporal scales, we propose TEP-Finder. In our experiment, TEP-Finder has been compared with NPM+ and DHAC+. The results demonstrate that TEP-Finder is better for capturing emerging phenomena and relationships among them.

## III. Definition of Emerging Phenomenon

In a long temporal phenotype dataset 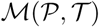, *T*_*i*_ is a time frame associated with experimental environments, treatments, and outcomes (*T*_*i*_ ∈ 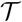), and *P*_*j*_ is an genotype to study (*P*_*j*_ ∈ 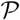), e.g., a gene knockout or a inbred line. The phenotype values of genotype *P*_*j*_ in time frame *T*_*i*_ are represented by a set of data points 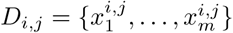. In the plant photosynthetic phenotyping experiment using DEPI [8], the phenotypes are mainly photosynthetic system II activity (Φ_*II*_), photoprotection (*q*_*ESV*_), and photoinhibition (*q*_*I*_). An emerging phenomenon *e*_*i*_ is defined as follows (see example in Figure 1).

### Definition III.1. Emerging Phenomenon.

Given the temporal phenotype data 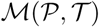, an emerging phenomenon *C*(*P_λ_*, *T_λ_*) is a group of genotypes *P_λ_* (*P_λ_* ⊆ 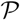) that exhibit coherent phenomena during continuous temporal range *T_λ_* (*T_λ_* ⊆ 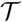), where |*P_λ_*| ≥ *K*_1_, |*T_λ_*| ≥ *K*_2_, and the percentage of significant phenotype values in *e* is greater than or equal to *K*_3_. *K*_1_, *K*_2_, and *K*_3_ are user specified thresholds.

Note that in an emerging phenomenon *C*(*P_λ_*, *T_λ_*), certain percentage of phenotype values of should be significantly different from the reference. The definition does not require all the phenotype values of *C*(*P_λ_*, *T_λ_*) to be significant because, when the environmental conditions vary dynamically, phenotype values often periodically switch between significance and insignificance (see Figure 1). Hence, it is more reasonable to require a certain portion but not all of the phenotype values to be significantly different from that of the reference. Since *A Priori* does not apply, new algorithms are needed for the identification of emerging phenomenon.

For a large-scale phenotyping experiment, the total number of identified emerging phenomena could be large. To better manage and use them, we construct an EP-DAG 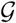 defined as:

### Definition III.2. Emerging Phenomenon DAG.

An emerging phenomenon DAG (EP-DAG) 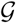 is a directed acyclic network (DAG), where each node in 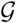 represents represents an emerging phenomenon *C*(*P_λ_*, *T_λ_*), and node *C*(*P*_*i*_, *T*_*j*_) is a descendent of node *C*(*P*_*h*_, *T*_*k*_) if and only if *P*_*i*_ ⊃ *P*_*h*_ and *T*_*j*_ ⊂ *T*_*k*_.

The outputted EP-DAG is available in the OBO format. It, once generated from data, can be visualized with Cytoscape [25] or OntoVisT [28]. It automatically supports emerging phenomenon search, phenotype relationship identification, and multiple phenotyping experiments comparison, leading to improved computational efficiency and succinct representation. To our knowledge, there is no existing work focused on the construction of EP-DAGs.

## IV. Methods

To systematically identify emerging phenomena in long-term phenotyping experiments and to examine the interactions between emerging phenomena and dynamic environments in a genetically diverse population, we introduce a new algorithm called **TEP-Finder**. TEP-Finder has two phases. First, it identifies seed phenomena in every time frame of a longitudinal phenomics dataset, where a time frame is a predefined minimal temporal range of any emerging phenomena. Second, by expanding each identified seed phenomena to longer time frames, TEP-Finder discovers emerging phenomena that appear and disappear subject to the change of environments or time. Multiple emerging phenomena are then merged, pruned, and connected to form a phenomenon network to facilitate phenotype search, comparison, and functional analysis. The diagram of the whole process is shown in Figure 2.

### A. TEP-Finder Phase 1. Identifying Seed Phenomenon

An emerging phenomenon *e* is considered as the phenotypes of multiple genotypes that have similar variation trends in a continuous time period. Biologically, such time period can be transient or can last for a relatively long time. To identify *e* with varying length, we consider a seed-based approach. Namely, we segment the whole experiment duration into multiple time frames, each being the minimal temporal range of any emerging phenomena. Then, at every time frame, we seek seed phenomena that are potentially extendable to a longer time period. The seed identification phase can be divided into four steps.

*1) Data Segmentation and Data Representation:* Given the temporal phenotype data 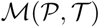, we segment 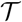 into separated time frames with a fixed length *m* using the sliding window approach. Here, the window width is the smallest temporal period of any emerging phenomenon (e.g., 30 minutes) that users can specify.

We adopt a meta-clustering approach to identify the relationships among all the tested genotypes in each time frame *T*_*i*_ [5], [18]. In the meta-clustering process, we repeatedly cluster the phenotype values of all the genotypes 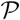 in *T*_*i*_ using non-parametric clustering with random anchor points [12]. The center of non-parametric clustering is a cloud-of-points representation. Since all the phenotype values are collected in a relatively short time, we examine the dependence among different phenotypes while ignoring the sequential order among the values and simply characterizing the phenotypes of genotype *P*_*j*_ in time frame *T*_*i*_ by the set of data points 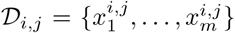, which we refer to as *cloud-of-points* representation.

*2) Phenotype Clustering:* Following the standard framework of mixture models, we assume that there are *K* different underlying distributions in time frame *T*_*i*_, where each distribution is introduced to capture a different “shape” of the cloud-of-points representation, and all the phenotype values observed in the cloud-of-points representation are drawn independently from one of the *K* distributions [10]. More specifically, let *f*_1_(·),…, *f*_*K*_(·) be the density functions for the *K* underlying distributions, and let *p*_1_,…, *p*_*K*_ be the prior probabilities for choosing each distribution. Then, for genotype *P*_*j*_ in time frame *T*_*i*_, the likelihood of observing the cloud-of-points representation 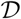_*i,j*_ is then given by

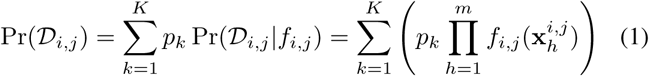

Following the framework of maximum likelihood estimation, we find the optimal density functions 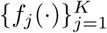 by solving the optimization problem

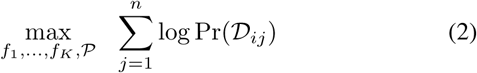

where Pr(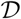_*ij*_) is given in Equation 1, and *n* is the total number of genotypes in 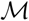.

This optimization problem can be effectively solved by employing NPM, a non-parametric clustering method for phenomics data analysis [12]. Based on the Nadaraya-Watson method for kernel density estimation [20], [24], [27] and following the framework of maximum likelihood estimation, NPM uses optimal density functions and applies a nonparametric clustering technique to group genotypes into the same cluster if their clouds-of-points share similar arbitrary shapes. The non-parametric approach avoids the parametric assumption of the underlying distribution so that NPM is suitable to model the nonlinear interactions among multiple phenotypes [12]. Since the clustering process is dependent on the initialization and the selection of anchor points, we repeat NPM multiple times to obtain all the meta-clustering results.

*3) Dynamic Phenotype Network Construction:* In this step, we construct a dynamic phenotype network 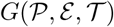, where 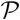 is the set of genotypes, 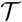 is the set of time frames, and *ε* = {*E*_1_, *E*_2_,…, *ε*_*k*_} represents edges in different time frames. In each time frame *T*_*i*_, we check whether any two genotypes *P*_*j*_ and *P*_*h*_ are frequently grouped into the same cluster in meta-clustering. If the co-occurrence is greater than a predefined threshold *K*_4_, we add edge 〈*P*_*j*_, *P*_*h*_〉 to *E*_*i*_. In the dynamic network *G*, while the nodes are identical, the edges vary over time, indicating emerging phenomena emerge and disappear with the change of time.

A running example is shown in Figure 3. In the example, 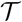 = {*T*_1_, *T*_2_, *T*_3_} and 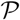 = {*A, B, C, D, E, F, G*}. Given the frequency of concurrence of every two genotypes in *T*_1_, *T*_2_ and *T*_3_ (the table on the left), and let *K*_4_ be 0.8, we identify all the edges (shaded blocks) and construct the dynamic network in the middle panel of Figure 3.

**Figure 3.**
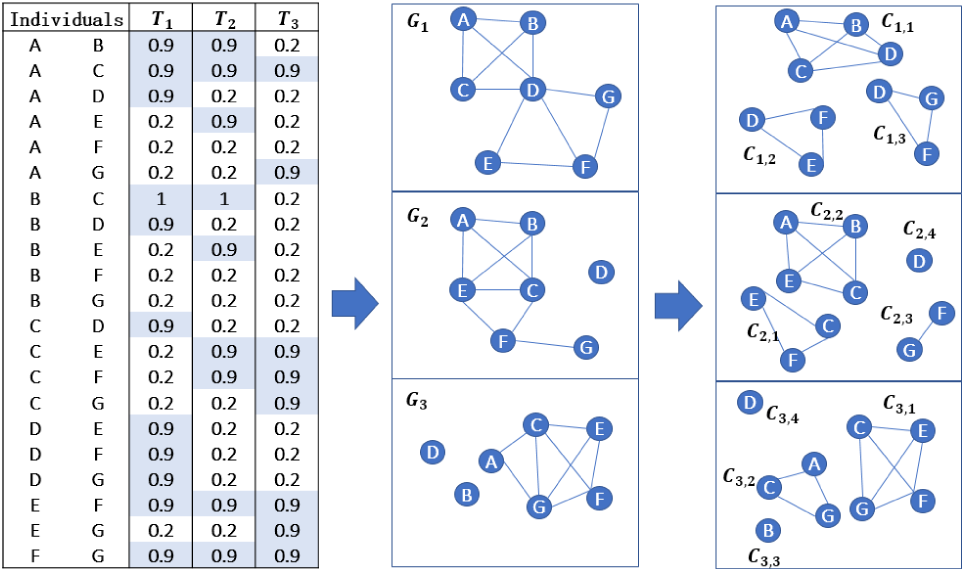
Illustrative example of the dynamic phenotype network construction. Values in the table represent the co-occurrence frequencies of any two genotypes being in the same cluster in three different time frames {*T*_1_, *T*_2_, *T*_3_}. We add edge 〈*P*_*j*_, *P*_*h*_〉 to a slice of the dynamic phenotype network if the corresponding co-occurrence frequency of genotypes *P*_*j*_ and *P*_*h*_ is greater than a threshold (shaded blocks). The middle panel shows the dynamic network, in each slice of which, the maximal cliques are displayed in the right panel.

*4) Seed Phenomena Identification:* We identify the seed phenomena by repeatedly applying a maximal clique based approach on every time frame of the dynamic network 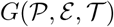. Clique is a special structure such that any two nodes in it are adjacent, implying a close relationship among all the nodes that belong to the same clique. A maximal clique is a clique that cannot be extended by including one more adjacent node, meaning it is not a subset of a larger clique.

More specifically, we adopt the Bron-Kerbosch algorithm to identify all the maximal cliques [4]. The basic form of the Bron-Kerbosch algorithm is the recursive backtracking that searches for all maximal cliques in a given network. Its performance has been further improved by defining a pivot vertex set, allowing it to backtrack more quickly in branches of the search that contain no maximal cliques [6], [30].

Let ℂ be the set of all the maximal cliques in the dynamic network 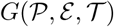, maximal clique *C*(*P*_*j*_, *T*_*i*_) ∈ ℂ defines a seed phenomenon with its genotype set being *P*_*j*_ and its time frame being *T*_*i*_. A running example is shown in the right panel of Figure 3. The maximal cliques in *T*_1_ are *C*_1,1_ = {*A, B, C, D*}, *C*_1,2_ = {*D, E, F*}, *C*_1,3_ = {*D, F, G*}.

### B. TEP-Finder Phase 2. Extending from Seeds to Emerging Phenomenon

After identifying all the seed phenomena in the minimal time frames, we extend them to longer time frames. This phase has four steps.

*1) Emerging Phenomenon Identification:* To identify emerging phenomena, the general idea is to combine seed phenomena in adjacent time frames. More specifically, for *C*(*P*_*j*_, *T*_*i*_), which is the *j*th seed phenomenon in time frame *T*_*i*_, we join it with every seed in time frame *T*_*i*+1_ *C*(*P*_*k*_, *T*_*i*+1_), resulting in *C*(*P*_*j,k*_, *T*_*i,i*+1_), where *P*_*j,k*_ represents the intersection of *P*_*j*_ and *P*_*k*_. Then, we determine whether *C*(*P*_*j,k*_, *T*_*i,i*+1_) is a new emerging phenomenon using the following rules developed based on the definition of the emerging phenomenon (see Definition III.2). The combination process will continue on the followed time frames (i.e. *T*_*i*+2_,…, *T*_*n*_), until all the seed phenomena are examined.

- Discard *C*(*P*_*j,k*_, *T*_*i,i*+1_) if |*P*_*j,k*_| < *K*_1_, is not an emerging phenomenon;
- Replace *C*(*P*_*j*_, *T*_*i*_) with *C*(*P*_*j,k*_, *T*_*i,i*+1_) if *P*_*j,k*_ = *P*_*j*_ ≥ *K*_1_.
- Replace *C*(*P*_*k*_, *T*_*i*+1_) with *C*(*P*_*j,k*_, *T*_*i,i*+1_) if *P*_*j,k*_ = *P*_*k*_ ≥ *K*_1_.
- Accept *C*(*P*_*j,k*_, *T*_*i,i*+1_) as a new emerging phenomenon if *S*_*j,k*_ ≠ *S*_*j*_ and *S*_*j,k*_ ≠ *S*_*k*_ and |*S*_*j,k*_| ≥ *K*_1_

Following the example of the dynamic phenotype network and maximal cliques in Figure 3, the emerging phenomenon identification procedure starting from *T*_1_ is shown in Figure 4. In Figure 4(a), one of the seed phenomena that start from *T*_1_ or *T*_2_ is *P*_2_ = *C*({*D, E, F*}, {*T*_1_}) and *P*_4_ = *C*({*C, E, F*}, {*T*_2_}) respectively. The join of *P*_2_ and *P*_4_ is *P*_9_ = *C*({*E, F*}, {*T*_1,2_}), which, according to Definition III.1, is saved as an emerging phenomenon in time frame *T*_1,2_. Similarly, for the other seeds in *T*_1_ and *T*_2_, we join them pair-wisely and save all the qualified emerging phenomena (see the blue colored notes in Figure 4(a)). Next, we join all the emerging phenomena in time frame *T*_1,2_ with the seeds in *T*_3_, resulting in the emerging phenomena in time frame *T*_1,2,3_. For example, *P*_18_ = *C*({*E, F*}, {*T*_1,2,3_}) is the result by joining *P*_9_ = *C*({*E, F*}, {*T*_1,2_}) and *P*_14_ = *C*({*C, E, F, G*}, {*T*_3_}). Note that *P*_18_ replaces *P*_9_ since they have the same genotypes and the time frame of *P*_18_ contains that of *P*_9_. Those who do not qualify the definition of emerging phenomenon are discarded (all the gray notes in Figure 4(a)).

**Figure 4.**
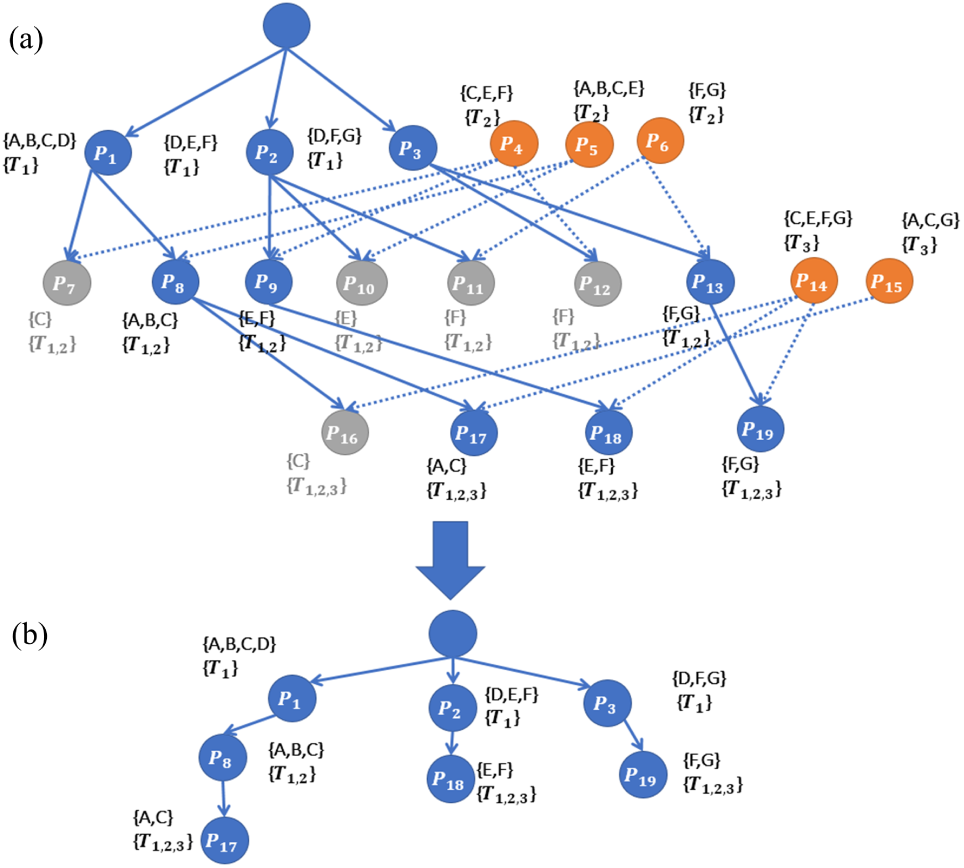
Illustrative example on extending a seed phenomenon to a longer time frame starting from the same time point. (a) The maximal cliques of *G*_1_ are at the first level. Then they are joined with the maximal cliques of *G*_2_ and *G*_3_ to generate longer emerging phenomena. (b) The result is pruned using the procedure introduced in Section IV-B1.

*2) Significance Test:* Given the temporal phenotype data 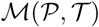, we compare the phenotype values of every genotype *P*_*i*_ with the reference using logged fold change, resulting in the relative phenotype values. The reference could be the wild-type in mutant experiments, the parental lines in recombinant inbred line experiments, or the average of all the genotypes in population experiments. Without losing generality, all the significant phenomena can be identified using a user given logged fold change threshold or with the computation of the false discovery rate. Other significance tests can also be applied for the same purpose. If the percentage of significant phenotype values of an emerging phenomenon is less than a user given threshold *K*_3_, the emerging phenomenon is discarded.

*3) EP-DAG Construction:* To model the complex relationships among all the emerging phenomena, we construct an EP-DAG 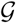. 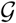 is a DAG with a virtual root node *P*_*root*_. We first connect all the emerging phenomena found in any individual time frame directly to *P*_*root*_ (see example in Figure 4(b)). Next, we add an edge pointing from every emerging phenomenon to another one if the latter is generated by joining the former with other ones and both of them start from the same time frame. For example, in Figure 4(b), an edge is pointing from *P*_8_ = *C*({*A, B, C*}, {*T*_1,2_}) to *P*_17_ = *C*({*A, C*}, {*T*_1,2,3_}). Finally, we add an edge pointing from one emerging phenomenon *C*(*P*_*j*_, *T*_*i*_) to another one *C*(*P*_*h*_, *T*_*k*_), if *P*_*j*_ ⊂ *P*_*h*_, *T*_*i*_ ⊃ *T*_*k*_, and *C*(*P*_*h*_, *T*_*k*_) is not a descendent of *C*(*P*_*j*_, *T*_*i*_). For example, we add edges pointing from *P*_5_ to *P*_8_, *P*_15_ to *P*_23_, and *P*_14_ to *P*_21_ (see the dotted edges in Figure 5(b)).

**Figure 5.**
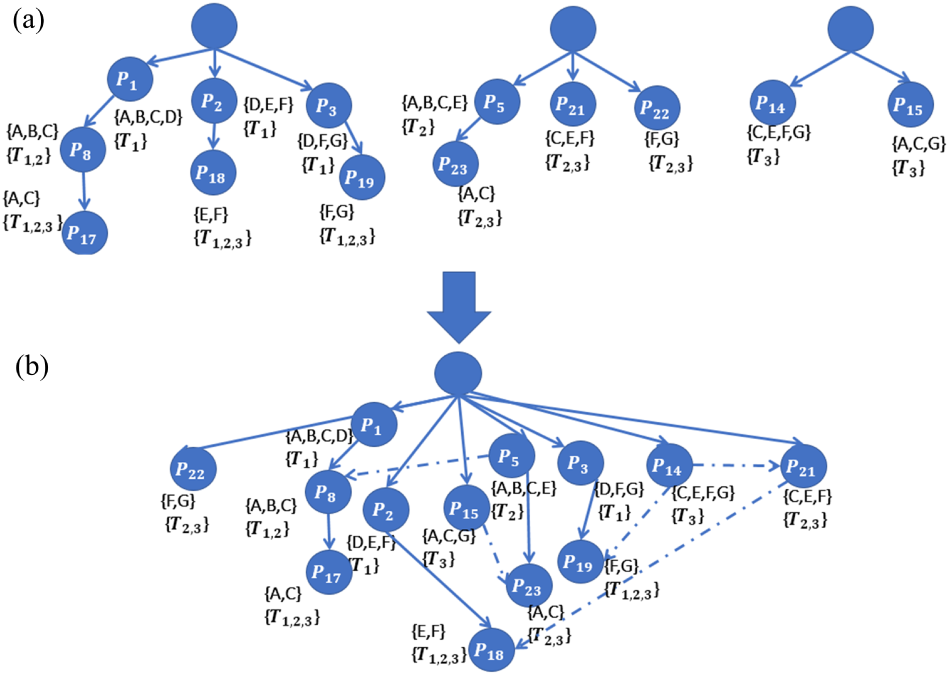
Illustrative example on EP-DAG construction. (a) Three sub-DAGs are built based on different starting time points. (b) All of them are merged into one DAG using the procedure introduced in Section IV-B3.

*4) EP-DAG Pruning:* Finally, to reduce the redundancy of the emerging phenomenon, we merge the highly overlapped emerging phenomena and remove emerging phenomena with insignificant phenotype values. Note that if an emerging phenomenon is discarded because it does not satisfy the user given thresholds (e.g., the percentage of significant values less than *K*_3_), its children will be redirect to its patent emerging phenomena. See examples in Figure 4(a,b). Mathematically, given two emerging phenomena *C*(*P*_*j*_, *T*_*i*_) and *C*(*P*_*h*_, *T*_*i*_) in the same time frame, if |*P*_*j*_ - *P*_*h*_| ≤ 1 and |*P*_*h*_ - *P*_*j*_| ≤ 1, we remove the two emerging phenomena and compose a new one called *C*(*P*_*j*_ ∪ *P*_*h*_, *T*_*i*_). Meanwhile, the edges connecting to *C*(*P*_*j*_, *T*_*i*_) and *C*(*P*_*h*_, *T*_*i*_) are redirected to *C*(*P*_*j*_ ∪ *P*_*h*_, *T*_*i*_).

## V. Results

### A. Data Description

For performance evaluation, we used the long temporal plant photosynthesis phenomics data in Gao et al [12]. The phenotyping experiment was carried out by testing 182 chloroplast-targeted single mutant lines (each with at least four biological replicates) of *A. thaliana* under dynamic environmental conditions using DEPI [8]. Three kinds of phenotypes, i.e., photosynthetic system II activity (Φ_*II*_), photoprotection (*q*_*ESV*_), and photoinhibition (*q*_*I*_) were collected at 112 time points. See experiment details in [8].

TEP-Finder was implemented with Python 2.7. The following parameters for TEP-Finder were used in the experiment: number of genotypes *K*_1_ = 5, number of time points *K*_2_ = 10, percentage of significant phenomena *K*_3_ = 0.5, number of time points per time frame 10, overlap rate between two adjacent time frames 90%; number of runs of clustering per time frame 100. The final results consist of 4,318 emerging phenomena and an EP-DAG with 7,789 edges.

### B. Methods to Compare

We compared TEP-Finder with NPM+ and DHAC+. The latter two are the methods modified from NPM and DHAC respectively (see Section II). The major difference in these methods locates on the process of seed phenomenon identification. More specifically, NPM+ consists of the following two steps. First, given the phenotype data 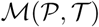, we call NPM once at every time frame to obtain the clustering results. Second, the clusters are used as the inputs to TEP-Finder phase two (see Section IV-B). In DHAC+, we first preprocess the phenomics data 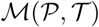 using the phase one of TEP-Finder, resulting in the dynamic phenotype network *G*. Second, DHAC is adopted to identify seed phenomena in *G* instead of searching for the maximal cliques. Finally, the seed phenomena are used as the inputs to TEP-Finder phase two (see Section IV-B). Note that the only difference between NPM+, DHAC+, and TEP-Finder is how the seed phenomena are identified. Comparing NPM+ and DHAC+ with TEP-Finder is critical because it can test whether our meta-clustering followed with maximal clique approach is appropriate to generate seeds, which form the basis for the identification of emerging phenomena.

### C. Performance Evaluation using GO Enrichment

An emerging phenomenon consists of a list of chloroplast-targeted single mutant lines that exhibit coherent and significant phenomena in a continuous time frame. It is expected that the knockout genes would be involved in the same biological process or have a similar molecular function. Therefore, we tested whether the knockout genes in the same emerging phenomenon are also enriched in Gene Ontology (GO). GO includes three categories: biological process, molecular function, and cellular component. Given a set of genes and their GO annotations, GO enrichment analysis identifies the overrepresented GO terms. In our experiment, data were downloaded from the GO website in March 2017, and clusterProfiler [34] was used for the enrichment test.

Figure 6(a) shows that the percentage of emerging phenomena at each level of the EP-DAG using GO biological process. Clearly, TEP-Finder is constantly better than DHAC+, *esp*. at deep levels of the EP-DAG. The high performance on deep levels is important because emerging phenomena at deep levels often represent abnormal photosynthetic behaviors in a relatively more extended time period. TEP-Finder and NPM+ have a similar trend, but TEP-Finder is still better than NPM+ on most of the cases. Specifically, the averaged percentage of the enriched emerging phenomena of TEP-Finder is 0.80, which is 0.70 for NMP+. Similar results are found on the GO enrichment test on the molecular function category. In general, the performance of TEP-Finder is higher than NPM+ and DHAC+ at each level of EP-DAG (Figure 6(b)). The averaged percentage of the enriched emerging phenomena of TEP-Finder is 0.66, while the values of NPM+ and DHAC+ are 0.56 and 0.43 respectively.

**Figure 6.**
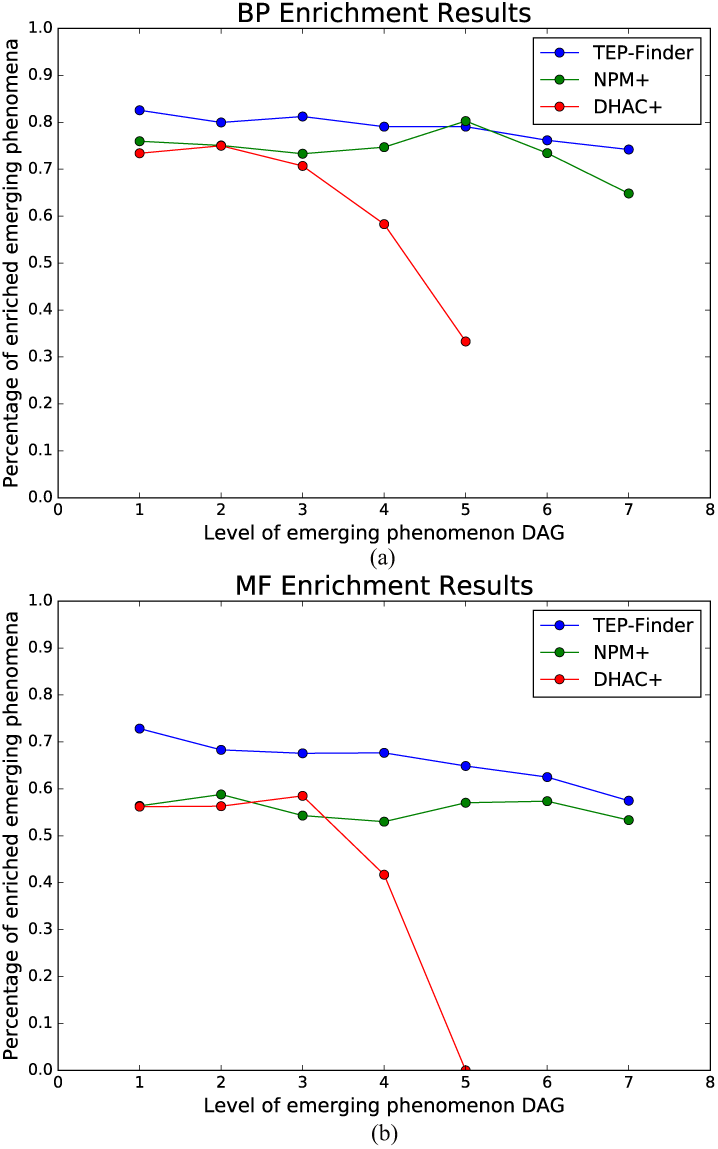
Evaluation of emerging phenomena using Gene Ontology enrichment on biological process (a) and molecular function (b). The x-axis represents the level of the EP-DAG. The y-axis represents the percentage of the emerging phenomena enriched in at least one GO term. Blue, green, and red represent the results of TEP-Finder, NPM+, and DHAC+.

While the first experiment shows that TEP-Finder has more enriched emerging phenomena than the other two, it is not clear whether the enriched GO terms are at a shallow or deep level of the GO. Therefore, in the second experiment, we compared the distribution of the enriched GO terms among the three methods. Since the first level of the EP-DAG is the virtual root node, the comparison was carried out at the second, third and fourth level. We only tested the first three valid levels of EP-DAG because, in the results of DHAC+, the number of emerging phenomena after three levels are too few to compare. Figure 7 shows the cumulative distribution of the GO biological process terms and the molecular function terms at the first three valid EP-DAG levels. It is constant that there are more deep-level enriched GO terms in TEP-Finder than the other two.

**Figure 7.**
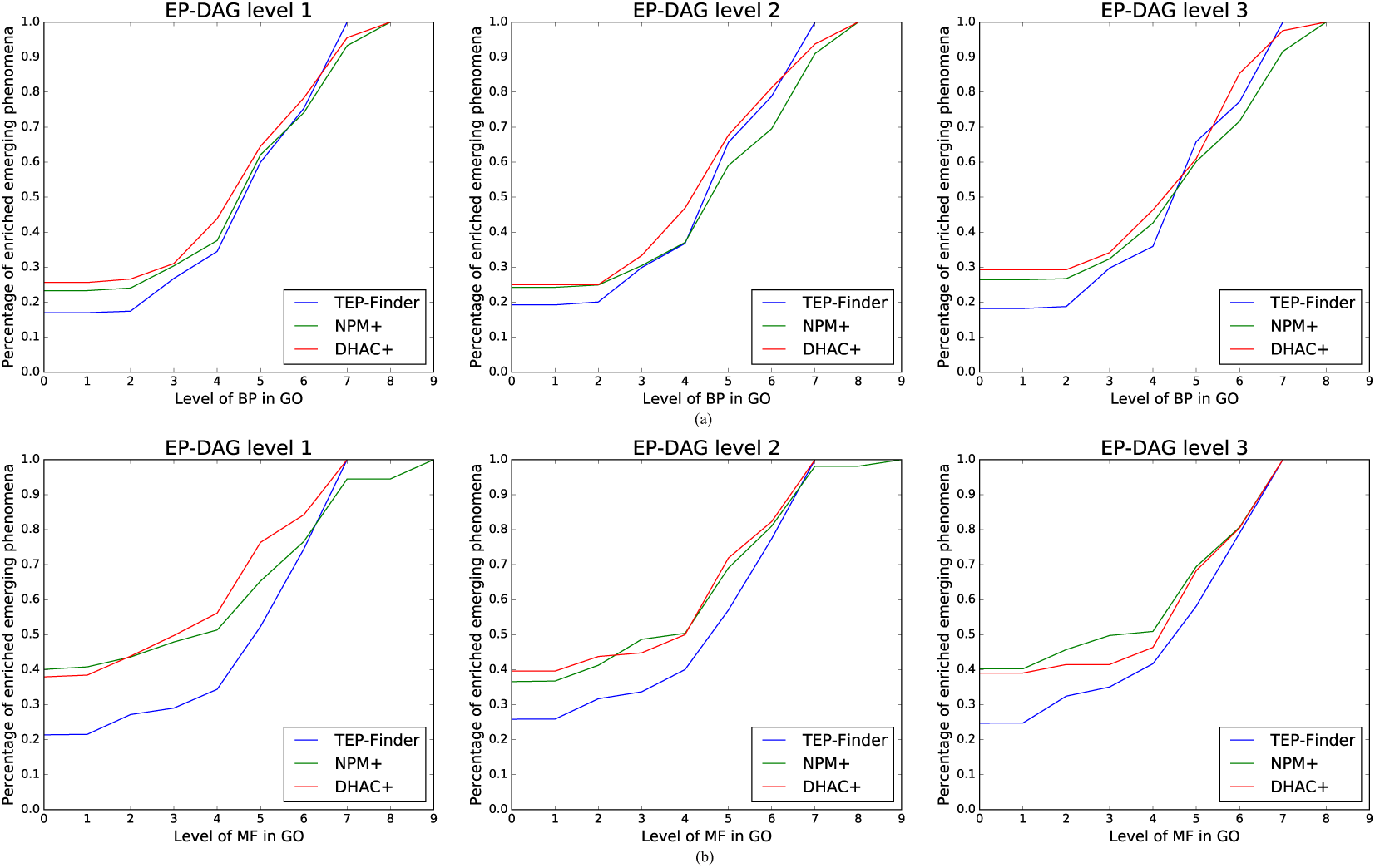
Cumulative distributions of identified emerging phenomena at different levels, which are enriched in Gene Ontology (GO) biological process category (a) and molecular function category (b). The x-axis represents the level of GO. The y-axis represents the percentage of emerging phenomena enriched at each GO level. The blue, green, and red line represent the result of TEP-Finder, NPM+, and DHAC+.

**Figure 8.**
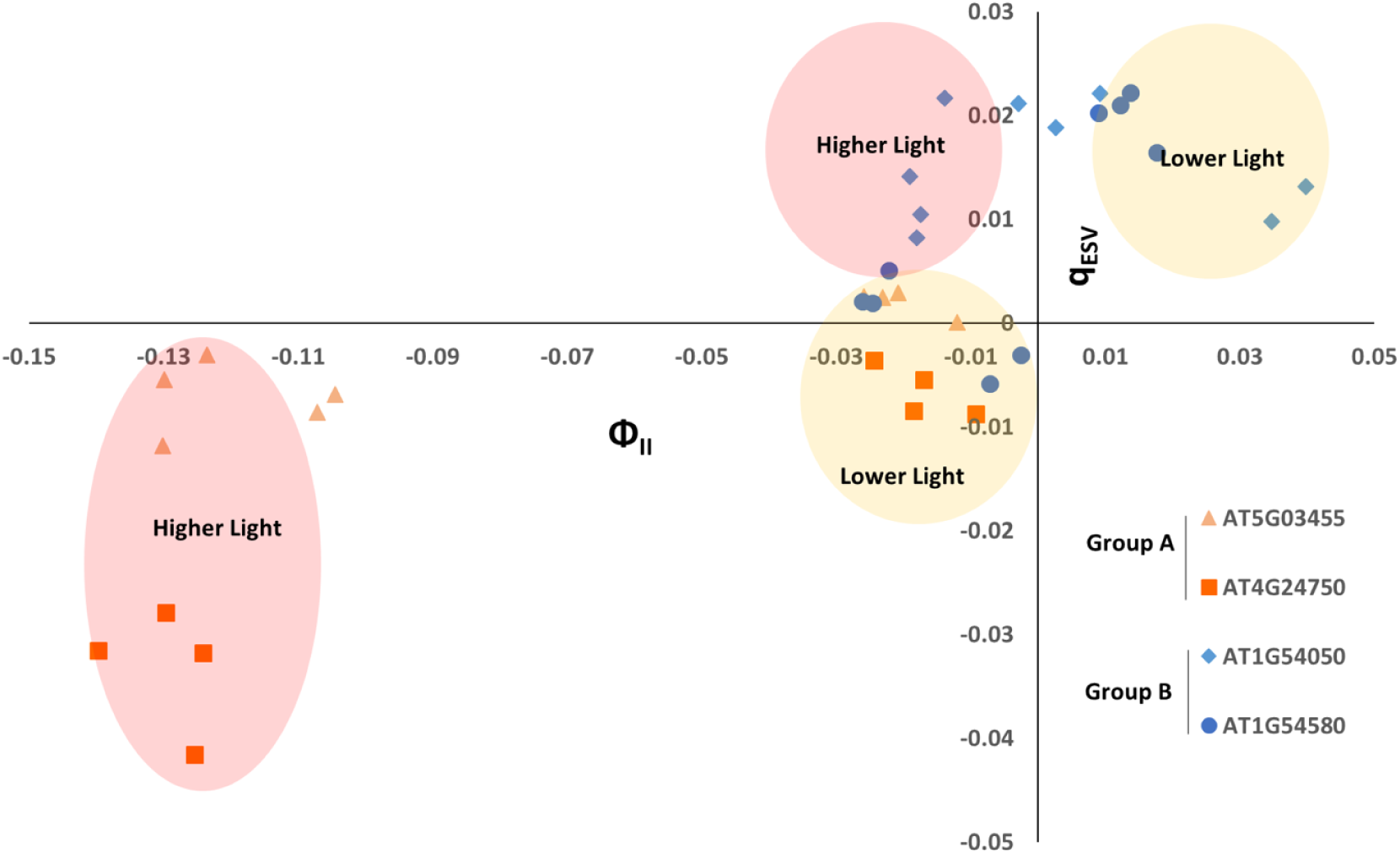
Two emerging phenomena found under strong fluctuating light conditions (between approximately 500*μmolm*^−2^*s*^−1^ (lower light) and 1000*μmolm*^−2^*s*^−1^ (higher light) four times repeated) have distinctively different photosynthetic phenotypes. Only two selected genotypes are shown for each group. In the first emerging phenomenon (group A, orange), plants have constantly low photoprotection yet the PS II activity decreased with the increased lights, indicating they are under stress. In the second one (group B, blue), less decrease of PS II activity and with high photoprotection as light increases, indicating they are well accommodated with the rapid changes of light.

### D. Performance Evaluation using Gene Association

Given an EP-DAG, gene-to-gene similarity can be calculated based on the topological structure of the DAG. We thereby test the correlation between EP-DAG based gene similarities with the GO molecular function based gene similarities. To calculate the gene-to-gene similarities based on the EP-DAG, we adopted the widely used Resnik method [23]. Specifically, given any two genes *g*_*i*_ and *g*_*j*_, we identify their least common ancestor term and calculate the gene-gene similarity using 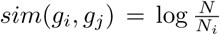, where *N* is the total number of genes in the ontology and *N*_*i*_ is the number of genes annotated to the lowest common ancestor. The GO-based gene similarities were calculated using a web service named InteGO2 [21]. All the similarities were normalized to the range of [0, 1].

Figure 9 shows the experimental results on the three networks constructed using TEP-Finder, NPM+, and DHAC+, perspectively. In general, there is a strong correlation between the gene-gene similarities based on TEP-Finder and based on the GO. Specifically, the *R*^2^ score of TEP-Finder is 0.89, significantly higher than that of the other two methods (i.e. 0.60 for NPM+ and 0.46 for DHAC+). This experiment suggests that the EP-DAG built by TEP-Finder is well organized.

**Figure 9.**
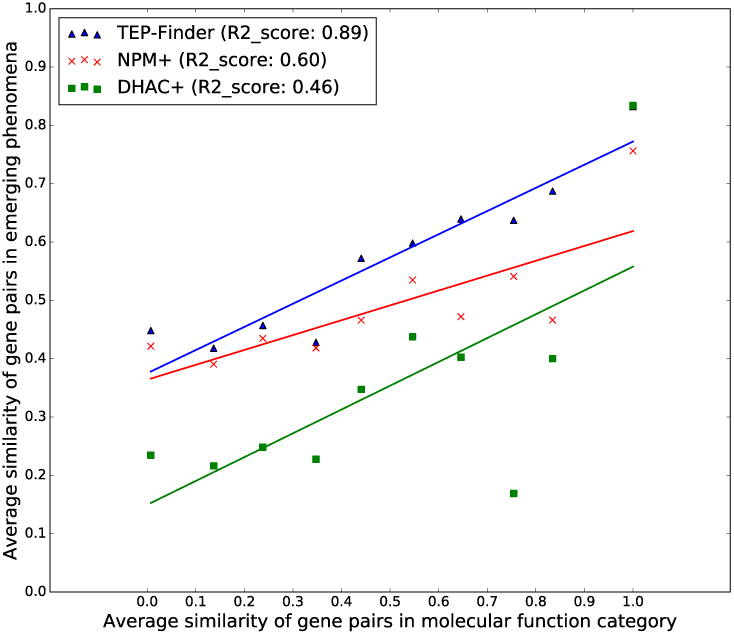
Comparing GO-based similarity with the EP-DAG based similarity. Gene pairs were clustered into 10 groups based on their GO-based similarities (x-axis), and for each group of gene pairs, we calculated the averaged EP-DAG based similarity (y-axis). Blue, green, and red represent the results of TEP-Finder, NPM+, and DHAC+.

### E. Biological Significance

Although we now have deep knowledge of the core processes of photosynthesis, the “ancillary components” essential for function in living cells under dynamic conditions are largely unexplored [35]. Intriguingly, these ancillary components probably evolved as plug-in functional modules to adapt the core processes to different conditions. Understanding their functions may allow us to combine these modules in different organisms, to achieve rapid improvements in the photosynthetic efficiency. The identification of the emerging phenomena of chloroplast-targeted single knockouts may enable systematic analysis of genotype-phenotype connections and provides a clue on the characterization of specific “ancillary processes” that support efficient photosynthesis. Note that many of the ancillary photosynthetic processes down-regulate the capture of light energy, preventing photodamage but at the cost of light-capture efficiency. From an evolutionary perspective, these processes can be viewed as balancing needs for energy and the avoidance of deleterious effects from photosynthesis.

We first analyzed the identified emerging phenomena from the gene evolution perspective. Since essential genes are often slow evolving compared with genes with nonlethal mutant phenotypes, the genes identified only in the emerging phenomena under fluctuating light varying conditions may evolve faster than those in the emerging phenomena under smooth light conditions. The ratio *Ka*/*Ks*, which measures the relative rates of synonymous and nonsynonymous substitutions at a particular site, is often used for the estimation of evolutionary rates [22]. In our experiment, the averaged *Ka*/*Ks* ratio of the 50 genes appeared only in the emerging phenomena under strong and smooth light conditions is 0.164, while the averaged *Ka*/*Ks* ratio of the 45 genes identified uniquely under fluctuating and strong light conditions is 0.192, significantly higher than the former (permutation test, p-value=0.013).

We then analyzed the emerging phenomena from the perspective of photosynthetic functionality. Two emerging phenomena (A and B) were categorized under the same strong fluctuating light conditions in the middle of the day (between 500*μmolm*^−2^*s*^−1^ and 1000*μmolm*^−2^*s*^−1^ four times repeated) due to distinctively different photosynthetic phenotypes (Figure 8 and S2, A, orange; B, blue). The emerging phenomenon A consists of mutant lines AT1G12250, AT1G80030, AT4G24750, and AT5G03455. They are sensitive to fluctuating light, showing large extent of decreases in PS II activity and decreases in *q*_*ESV*_ (photoprotection) under high light intensity compared to the low light. Mutant lines in emerging phenomenon B (AT1G14590, AT1G54580, AT2G40400, AT3G10470, AT4G31560, AT5G03455, AT5G39830) have less extent of decreases in PS II activity with higher *q*_*ESV*_ indicating less sensitivity to the fluctuating light. As the important genes responsive to dynamic light conditions, sensitivity of mutant would be increased. Thus, for mutant lines that are shown a sensitive phenotype under the conditions, it indicates that the mutated genes are responsible for maintaining robust photosynthesis under the stress conditions. Hence, we hypothesize that the genes in A may contribute to photoprotection in response to natural light dynamics (see the selected samples in Figure 8). According to the GO, these genes are involved in arsenate reductase activity and the photosynthesis-related biological processes, including arsenate reductase activity and oxidation-reduction process. Most of them are related to cellular redox balance, which are important for regulation photosynthesis, yet mode of function of found genes in this study is still partly remained elusive [3], [14]. This analysis may provide new insights and open the new possibility to understand how plants are adapted dynamic conditions. Mutant lines in B stay high *q*_*E*_ and minor decrease in PS II activity in the high light indicating those mutants are less sensitive to dynamic light conditions. It shows that the mutate genes in B are less likely responsible to adapt fluctuating light conditions. A functional analysis based on GO shows that these genes are involved in cell cycle, cell division and protein complex oligomerization, and protein folding fatty acid biosynthetic process cytochrome *b*_6_*f* complex assembly. Also, the averaged *Ka*/*Ks* ratio of A and B is 0.24 and 0.22 perspectively, which is significantly higher than that of randomly selected chloroplast-targeted genes (permutation test, p-value=0.024 and 0.013).

The biological analysis demonstrates that accurately identifying emerging phenomena from plant phenotyping data may be valuable towards the characterization of specific ancillary processes that support efficient photosynthesis.

## VI. Discussion and Conclusion

Comprehensive analysis of emerging phenomena is required to improve our understanding of the quantitative variation of complex phenotypes and to attribute gene functions [11]. However, unlike frequent patterns, emerging phenomena may re-occur frequently or may appear only once during an experimental period, depending on the experimental design. TEP-Finder is the first tool towards capturing the emerging phenomena in large-scale longitudinal phenotyping experiments, leading to the identification of the minimum set of distinct actors needed to produce an undefined, complex aggregate phenotypic trait. Particularly, TEP-Finder can identify emerging phenomena in different temporal scales from the data and also can construct a directed acyclic network (EP-DAG) for better data management. The Gene Ontology and gene-gene association based performance evaluation show that TEP-Finder is better than the existing tools regarding biological significance.

An important component of TEP-Finder is the meta-clustering that repeatedly calls NPM with random anchor points for kernel density estimation. We tested whether the meta-clustering approach can lead to more robust results. Specifically, given the same input, we ran TEP-Finder and NPM+ three times and calculated the differences between the results of the three runs. Figure S1 shows the average number of genes per emerging phenomenon (indicated by circle area) at each level of the EP-DAG. In Figure S1(a), the three runs of TEP-Finder are similar to each other, indicated by the highly overlapped circles, whereas the three runs of NPM+, as shown in Figure S1(b), are distinctively different. In summary, the adoption of the meta-clustering approach ensures TEP-Finder to be robust enough for emerging phenomenon mining.

A key parameter in capturing emerging phenomena is *K*_3_, the percentage of significant phenotype values. Unlike *K*_1_ and *K*_2_ that define the dimension of an emerging phenomenon, which is common in pattern recognition, *K*_3_ is difficult to specify. Here we fixed *K*_1_ and *K*_2_ and varied *K*_3_ to explore rules for choosing *K*_3_. Table S1 indicates that with the increase of *K*_3_, the EP-DAG becomes more concise (more shallow and has less amount of nodes), and the majority of the removed nodes are intermediate nodes. It suggests that to choose an optimal *K*_3_, we can start with a high value and then gradually reduce it. At the same time, we should check whether the leaf nodes (which has the longest time frames) captures long-term patterns. As a future work, we will develop new algorithms to automatically optimize the parameters of TEP-Finder.

